# The adhesion molecules β7 integrin and L-selectin contribute to cholestatic liver disease in male mice

**DOI:** 10.1101/2025.09.24.678237

**Authors:** Agnes Seeger, Anshu Babbar, Sreepradha Eswaran, Maria Adams, Moritz Muschaweck, Nicole Treichel, Thomas Clavel, Guo Yin, Adrien Guillot, Kerstin Scheuer, Sabine Hamm, Jakob-David Adam, Norbert Wagner, Angela Schippers

## Abstract

**Background **& Aims**:** Primary sclerosing cholangitis (PSC) is a long-term progressive disease, often occurring in conjunction with inflammatory bowel disease (IBD). Dysregulated immune cell migration and gut microbiota alterations are implicated in disease progression. Livers of PSC patients show upregulation of the endothelial ligand mucosal addressin cell-adhesion molecule-1 (MAdCAM-1). Here we examined the role of the leukocytic adhesion molecules (AM) β7 integrin and L-selectin, both binding partners of MAdCAM-1, in an experimental mouse model resembling aspects of human PSC.

**Methods:** Wild type (WT), β7 integrin-deficient (β7^-/-^), L-selectin-deficient (L-sel^-/-^), and L-selectin/β7 integrin double-deficient (L-sel^-/-^/β7^-/-^) male mice were compared in the model of 3,5-diethoxycarbonyl-1,4-dihydrocollidine (DDC)-induced cholangiopathy. The extent of pathology was evaluated by serum parameters, histology, flow cytometry, and expression of inflammatory mediators. Fecal microbiota changes were assessed by 16S rRNA amplicon sequencing and intestinal permeability was measured by FITC-dextran assay.

**Results:** AM-deficient mice were markedly protected from DDC-induced cholangiopathy. Hepatic immune cell populations of AM-deficient mice differed significantly from those of WT mice. Adoptively transferred β7^-/-^ CD8^+^ T cells caused significantly less liver damage than CD8^+^ WT T cells in DDC-treated β7^-/-^ mice. DDC-feeding caused substantial changes in fecal microbiota profiles, which differed between the mouse strains, and a strong increase in intestinal permeability that was significantly lower in β7^-/-^ than WT mice.

**Conclusions:** β7 integrin and L-selectin contribute to DDC-induced cholangiopathy with β7 integrin-expressing CD8^+^ T cells playing a crucial role in promoting pathogenesis. In addition, AM-expressing immune cells may contribute to the inflammatory process by causing unfavorable gut microbiota shifts and destabilizing the gut barrier.

**Synopsis:** β7 integrin- and/or L-selectin-deficient mice are less susceptible to 3,5-diethoxycarbonyl-1,4-dihydrocollidine (DDC)-induced experimental cholangiopathy. β7 integrin-expressing CD8^+^ T cells contribute to DDC-induced pathogenesis. β7 integrin promotes DDC-induced gut barrier dysfunction and microbiota shifts, potentially exacerbating hepatic injury.

## Introduction

Primary sclerosing cholangitis (PSC) is an immune-mediated progressive liver disease which is characterized by inflammation of the intra- and extrahepatic bile ducts. It causes peribiliary strictures, dilation of bile ducts, and fibrosis, resulting in impaired bile flow and consecutive liver damage, eventually leading to biliary cirrhosis and end-stage liver disease [1]. Around 1/3 of patients also develop cholangiocarcinoma in the course of the disease [2]. Liver transplantation remains the only treatment option to date [3]. More than 70 % of PSC patients suffer in addition from inflammatory bowel disease (IBD). The two main forms of IBD are Crohn’s disease and ulcerative colitis [4], with most PSC patients suffering from the latter [5]. The etiology of PSC is currently unknown, but a number of hypotheses for its development and progression are under consideration [6–8]. Based on the identification of predisposing genetic factors and the reduced risk of smokers developing PSC, an interaction of genetic variations and environmental influences is assumed [8, 9]. The inflammatory processes in PSC are characterized by increased infiltration of pro-inflammatory immune cells, suggesting the involvement of dysregulated cell trafficking in the pathogenesis. Given the close association of PSC and IBD, the hypothesis has been put forward that gut-primed immune cells are responsible for both intestinal and liver inflammation. It is assumed that these pro-inflammatory cells are able to infiltrate both organs due to the shared expression of specific homing signals, including chemokines, adhesion molecules and their respective ligands [10]. This hypothesis is supported by the detection of intestinal and liver T cells of common clonal origin in tissue of patients with IBD and PSC [11].

Further support for the above hypothesis is given by the fact that expression of the endothelial ligand mucosal addressin cell adhesion molecule-1 (MAdCAM-1), which in homeostasis is almost exclusively found in the gut, is upregulated on the hepatic sinusoidal endothelium of patients with chronic liver disease, including PSC patients [12, 13]. MAdCAM-1 is the major ligand for the adhesion molecule α4β7 integrin but binds also to L-selectin [14]. L-selectin has already been shown to play a role in driving metabolic dysfunction-associated steatotic liver disease (MASLD) [15]. β7 integrin forms heterodimers with either the α4 or the αE (CD103) subunit and is expressed on all immune cells except neutrophils [14]. While α4β7 integrin plays a key role in mediating immune cell homing to the intestine and mesenteric lymph nodes via binding to MAdCAM-1, αEβ7 integrin facilitates their retention by interaction with the epithelial ligand E-cadherin [16–18]. MAdCAM-1/α4β7 integrin interactions are well known to promote IBD and in accordance with the hypothesis presented above, α4β7 integrin-expressing lymphocytes have also been detected in the inflamed livers of PSC patients [13, 19].

In addition to disease-relevant cell trafficking events, other processes in the reciprocal gut-liver axis might be disturbed and are discussed in the context of PSC development. According to the leaky gut hypothesis, PSC is associated with breaching of the intestinal epithelial barrier which leads to increased intestinal permeability, allowing bacterial translocation into the liver. There, activation of the innate immune system by bacterial-derived products causes a pro-inflammatory milieu by inflammasome activation and pro-inflammatory cytokine production. When unresolved, prolonged inflammation has detrimental effects on parenchymal and non-parenchymal liver cells, ultimately causing liver fibrosis and liver failure [8, 20, 21]. The study of Nakamoto et al. provided first proof of this hypothesis in patients with PSC and ulcerative colitis [22]. It clearly demonstrated a pathogenic role of *Klebsiella pneumonia*, a bacterium which is increased in PSC patients and directly disrupts the intestinal epithelial barrier, thereby allowing bacterial translocation and subsequent liver inflammation.

This and other findings also indicate a direct involvement of the intestinal microbiota in the pathogenesis of PSC. A number of studies have shown gut microbiota changes in PSC patients [23–25] and some antibiotics have been found to reduce levels of PSC-related markers of liver injury [26]. Most interestingly, a connection between microbiota and retinoic acid metabolism was found, which in turn might contribute to the expression of homing molecules on immune cells, thereby influencing the infiltration of pro-inflammatory immune cells into gut and liver [27–31].

Although gut-derived cell trafficking pathways might represent promising targets for future therapeutic interventions in PSC they are still insufficiently explored. Our objective, therefore, was to study the role of β7 integrin and L-selectin expressing cells and their migration behavior in a mouse model of cholangiopathy. Feeding 3,5-diethoxycarbonyl-1,4-dihydrocollidine (DDC) is a widely used model for xenobiotic-induced cholangiopathies in mice. It causes increased biliary porphyrin deposition with formation of intra-ductal plugs, ductular reaction, pericholangitis accompanied by onion skin-type periductal fibrosis, immune cell accumulation, and subsequent development of biliary fibrosis, thereby producing many histological features of human PSC [32]. Utilizing this model, we compared the severity of cholangiopathy in wild type (WT), β7-integrin-deficient (β7^-/-^), L-selectin-deficient (L-sel^-/-^), and L-selectin/β7-integrin double-deficient (L-sel^-/-^/β7^-/-^) mice and were able to highlight the importance of these adhesion molecules in promoting the pathogenesis of cholangitis.

## Results

### Adhesion molecule-deficient mice are protected from DDC-induced liver injury, fibrosis and inflammation

To determine the role of the MAdCAM-1 receptors α4β7 integrin and L-selectin in the pathogenesis of cholangiopathy, we compared liver injury in male WT, β7 integrin-deficient (β7^-/-^), L-selectin-deficient (L-sel^-/-^) and L-selectin/β7 integrin double-deficient (L-sel^-/-^/β7^-/-^) mice after 4 weeks of 0.1% DDC feeding. First, we evaluated the suitability of this model for our purposes by checking whether the DDC-induced pathogenesis in WT mice is accompanied by an upregulation in the expression of adhesion molecules in the liver pointing to their contribution to the inflammatory process in this model. As expected, we detected significantly increased *Icam-1*, *Madcam-1* and *Vcam-1* mRNA levels in liver tissue homogenates of DDC-treated WT mice in comparison to the respective untreated controls (Figure 1A). DDC treatment induced a substantial weight loss in WT mice, which was less pronounced in all adhesion molecule-deficient mouse strains, with the least weight loss in L-selectin/β7 integrin double-deficient mice (Figure 1B). Levels of serum alkaline phosphatase and of direct bilirubin, which are markers of cholestasis and biliary injury, respectively, were markedly elevated in all mice after DDC feeding, but were substantially lower in β7 integrin-deficient and reduced even further in L-selectin/β7 integrin double-deficient mice. (Figure 1C, D). Previous studies showed that levels of serum aminotransferases, which are indicators of hepatocyte injury, increase just one week after starting DDC treatment but decline again afterwards [33, 34]. Serum levels of both aspartate aminotransferase (AST) and alanine aminotransferase (ALT) were found to be markedly increased in all our DDC-treated mice compared to untreated controls, irrespective of the genotype. However, the increase in serum enzymes of L-selectin/β7 integrin double-deficient mice after 2 weeks of DDC treatment was considerably lower than in all other mouse groups (Figure 1E, F). In addition, an amelioration of the DDC-induced liver tissue pathology was observed in β7 integrin-deficient and was even more pronounced in L-selectin/β7 integrin double-deficient mice, as reflected in a significant decrease in porphyrin deposition (Figure 1G, H). Moreover, Sirius red staining of liver sections confirmed extensive liver fibrosis in WT mice post DDC-feeding, whereas all similarly treated mutant mice were significantly less affected (Figure 1I, J).

**Figure 1:**
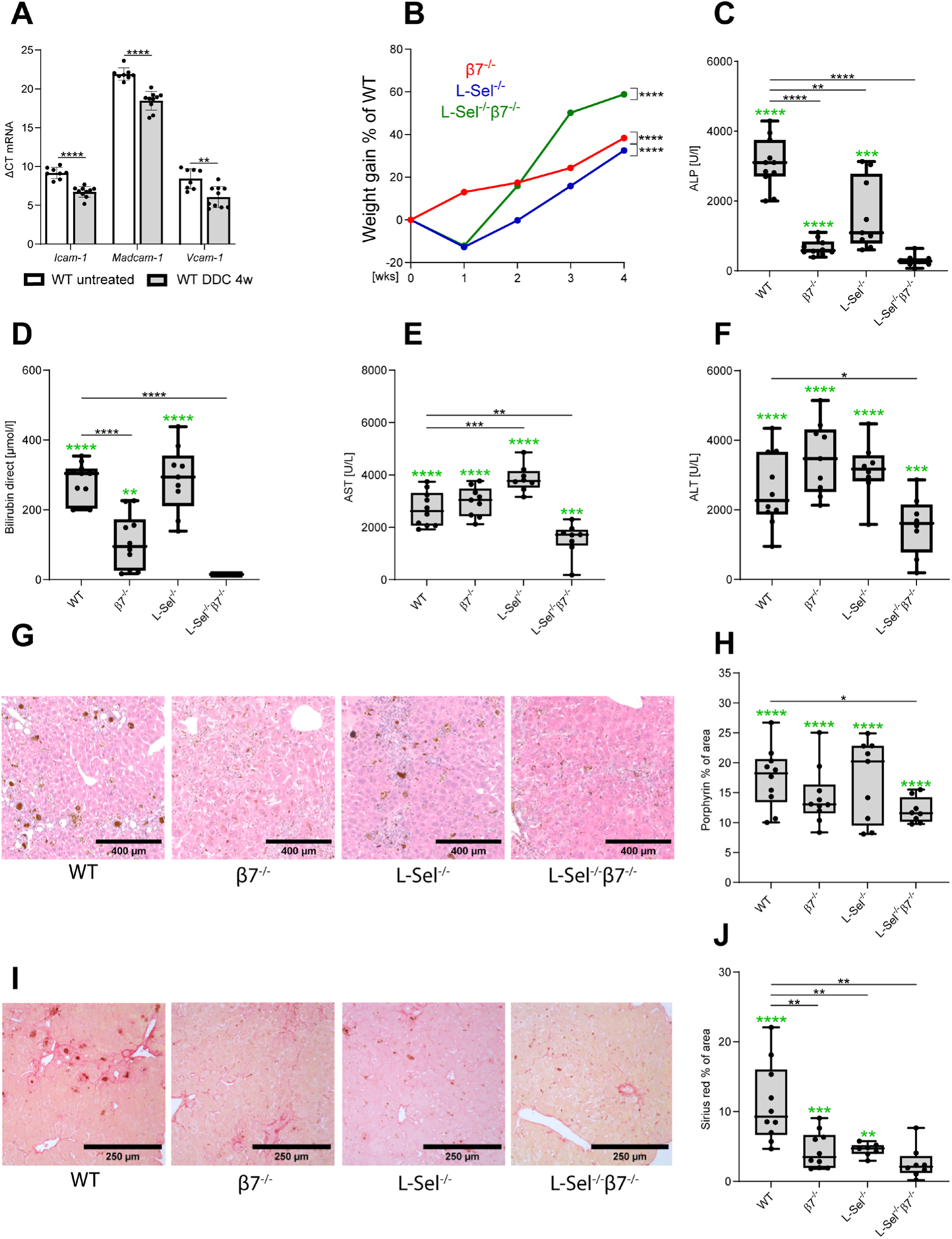
Adhesion molecule-deficiency ameliorates DDC-induced liver pathology. Serum and liver specimens were sampled from untreated, 2 week and 4 week DDC-treated mice. WT mice (untreated: n = 9, DDC-treated: n = 10-11), β7 integrin-deficient mice (β7^-/-^) (untreated: n = 9, DDC-treated: n = 9-10), L-selectin-deficient mice (L-sel^-/-^) (untreated: n = 10, DDC-treated: n = 8-9) and L-selectin/β7 integrin double-deficient mice (L-sel^-/-^/β7^-/-^) (untreated: n = 8, DDC-treated: n =8) (A) Analysis of the mRNA of the indicated adhesion molecules in liver tissue from untreated and 4 week DDC-treated WT mice. For quantification, values are expressed as delta (Δ) ct values between the genes of interest and the housekeeping gene *β actin*. (B) Percentage of weight gain from the indicated adhesion molecule-deficient mouse strains (measured weekly) related to similarly treated WT mice. (C-J) Values and images of DDC-treated mice only are shown here. Statistical significance of deviation from untreated control mice for each mouse strain is shown in panels C, D, E, F, H, and J as green asterisks (upregulation compared to control) above each group. (C) Quantification of alkaline phosphatase (ALP) and (D) quantification of direct bilirubin in serum after 4 weeks of DDC treatment. (E) Quantification of serum aspartate aminotransferase (AST) and (F) serum alanine aminotransferase (ALT) of the indicated mouse strains after 2 weeks of DDC treatment. (G) Representative photomicrographs of H&E stained liver sections from the indicated mouse strains after 4 weeks of DDC treatment at original magnification x10. (H) Quantification of porphyrin deposition in H&E stained liver sections from the indicated mouse strains after 4 weeks of DDC treatment shown as percentage of the section area. (I) Representative photomicrographs of Sirius red stained liver sections from the indicated mouse strains after 4 weeks of DDC treatment at original magnification x10. (J) Quantification of Sirius red stained collagen on liver sections from the indicated mouse strains after 4 weeks of DDC treatment shown as percentage of the section area. Statistical significance was calculated by the unpaired t test. Values are represented as mean ±SD and each dot (in A, C, D, E, F, H, and J) represents data from one mouse (*p < 0.05, ** p < 0.01, *** p < 0.001, ****p < 0.0001).

Next, we investigated whether the observed amelioration in clinical and histological parameters in adhesion molecule-deficient mice was also reflected in changes in expression levels of genes that are known to be involved in liver fibrosis, proliferation, and inflammation. In the livers of all mice, DDC-treatment resulted in an increased expression of the multidrug-resistance gene 1 (*Mdr1*) which is a membrane transport protein mediating efflux of a number of xenobiotics and toxins [35] (Figure 2A). In addition, the organic solute transporter beta (*Ost-β*), an alternative system for bile transport, and *Cyp2b10*, representing a nuclear receptor regulating bile acid detoxification downstream of the constitutive androstane receptor (CAR) [36], were substantially increased by DDC treatment in all mice (Figure 2B and C). Moreover, hepatic mRNA expression of monocyte chemoattractant protein-1 (*Mcp-1*), a key chemokine that regulates the migration of monocytes, was significantly upregulated in mice of all genotypes (Figure 2D). However, DDC-fed adhesion molecule-deficient mice exhibited a markedly reduced expression of *Ost-β*, *Cyp2b10* and *Mcp-1* when compared with similarly treated WT mice. By contrast, there was no significant difference in the DDC-induced enhancement of *Mdr-1* expression between WT and mutant mice (Figure 2A-D). Increased hepatic expression of the adhesion molecules VCAM-1, ICAM-1 and MAdCAM-1 is a hallmark of PSC pathogenesis [37]. Upon DDC feeding all mice showed a significant enhancement in hepatic mRNA levels for *Icam-1*, *Madcam-1* and *Vcam-1*. Interestingly, the increase in *Vcam-1* expression was significantly reduced in DDC-treated L-selectin-deficient and L-selectin/β7 integrin double-deficient mice, whereas β7 integrin-deficient mice exhibited a trend towards increased mRNA levels for *Madcam-1* and *Vcam-1* in comparison to similarly treated WT mice (Figure 2E-G).

**Figure 2:**
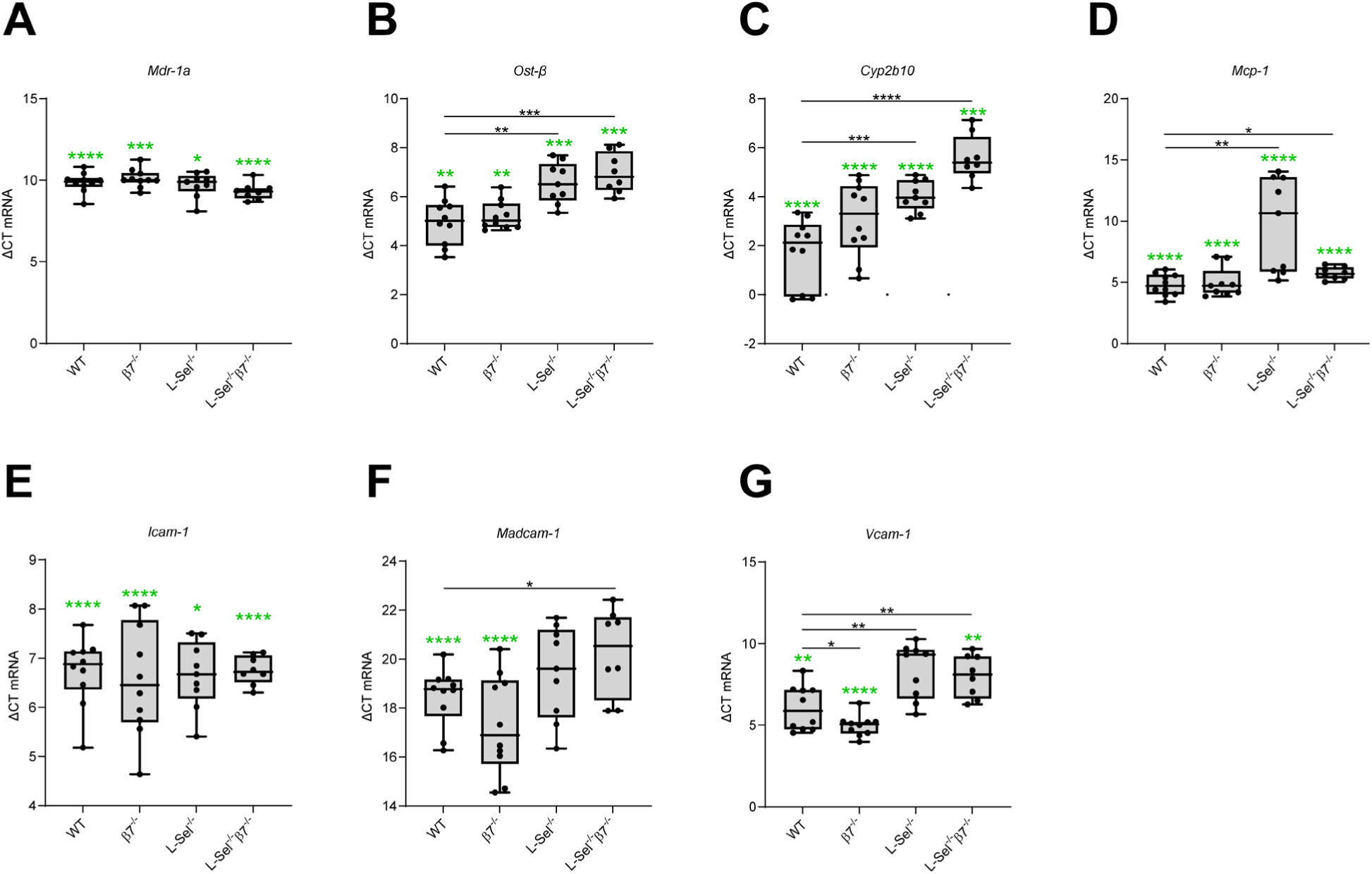
Adhesion molecule-deficiency results in decreased production of DDC-induced proinflammatory and profibrotic genes. Analysis of mRNA from liver specimens that were sampled from the indicated untreated and 4 week DDC-treated mouse strains by RT-qPCR. Values from 4 week DDC-treated mice only are shown here. Statistical significance of deviation from untreated control mice for each mouse strain is shown as green asterisks (upregulation compared to control) above each group. WT mice (untreated: n = 8, DDC-treated: n = 10), β7 integrin-deficient mice (β7^-/-^) (untreated: n = 9, DDC-treated: n = 10), L-selectin-deficient (L-sel^-/-^) (untreated: n = 10, DDC-treated: n = 9) and L-selectin/β7 integrin double-deficient (L-sel^-/-^/β7^-/-^) (untreated: n = 7, DDC-treated: n =8). For quantification, values are expressed as delta (Δ) ct values between the genes of interest and the housekeeping gene *β actin*. (A) *Mdr-1a* (*multidrug-resistance gene 1a*) (B) *Ost-β* (*organic solute transporter beta*), (C*) Cyp2b10*, (D) *Mcp-1* (*monocyte chemoattractant protein -1*), (E) *Icam-1* (*intercellular adhesion molecule-1*), (F) *Madcam-1* (*mucosal addressin cell-adhesion molecule-1*) (G) *Vcam-1* (*vascular cell adhesion protein-1*). Statistical significance of deviation from healthy control mice for each mouse strain is shown as green asterisks (upregulation compared to control) above each group. Statistical significance was calculated by the unpaired t test. Values are represented as mean ±SD and each dot represents data from one mouse (*p < 0.05, ** p < 0.01, ****p < 0.0001).

To establish whether the diminished inflammatory response following DDC treatment can be explained by reduced recruitment of leukocytes in the liver, we investigated the effects of a lack of adhesion molecules on the composition of the immune cell populations in the liver by flow cytometry. DDC feeding induced a marked influx of monocytes/macrophages (CD45^+^, CD11c^-^, CD11b^+^, Ly6C-6G^-^, F4/80^+^), Gr-1^+^ cells (CD45^+^, CD11c^-^, CD11b^+^, Ly6C_6G^+^), and CD8^+^ T cells into the livers of all mouse strains, whereas the number of CD4^+^ T cells changed only slightly (Figure 3A-D). Compared to livers of DDC-treated WT mice, β7 integrin- and L-selectin/β7 integrin double-deficient mice had significantly fewer macrophages, L-selectin/β7 integrin double-deficient mice fewer neutrophils, β7 integrin- and L-selectin-deficient mice fewer CD4^+^ T cells. The number of CD8^+^ T cells was most increased in adhesion molecule-deficient mice (Figure 3A-D). Since CD8^+^ T cells reportedly play an important role in PSC pathogenesis [38, 39] the presence of newly immigrated CD8^+^ T cells was also analyzed by the more sophisticated technology of multiplex immunofluorescence staining on liver sections. Here, too, it is clearly visible that DDC-treatment caused a strong migration of CD8^+^ T cells into the liver. In comparison to WT significantly more CD8^+^ T cells have migrated into the livers of β7 integrin- and L-selectin/β7 integrin double-deficient mice (Figure 3E and F). To determine whether the absence of β7 integrin affects CD8^+^ T cell migration into the liver, we compared the migration efficiency of β7 integrin-deficient and WT leukocytes by short-term migration assays in WT recipients. As expected, migration of β7 integrin-deficient CD8^+^ T cells into the Peyer’s patches was severely impaired, however, significantly more β7 integrin-deficient than WT CD8^+^ T cells have reached the liver. (Figure 3G).

**Figure 3:**
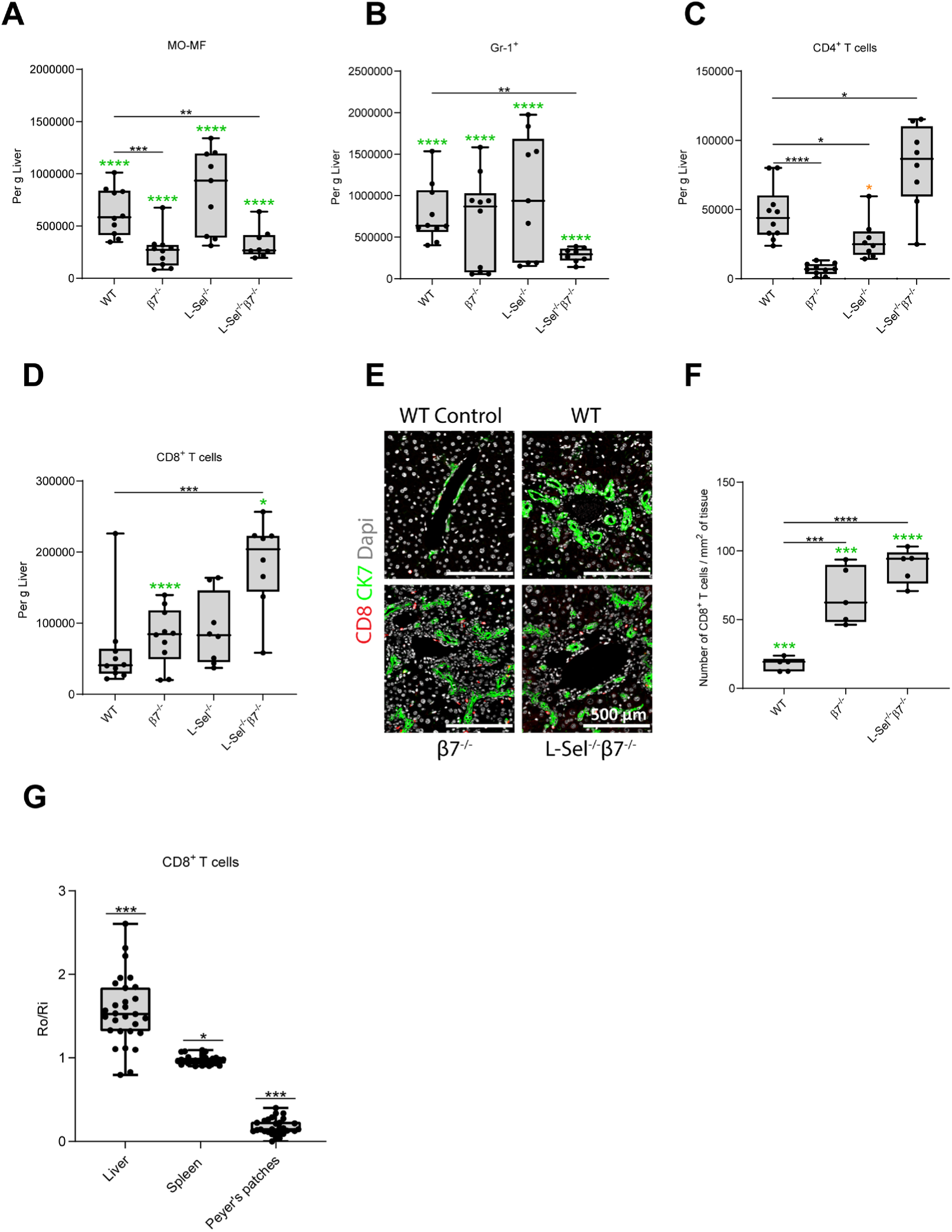
Absence of adhesion molecules alters hepatic immune cell populations in DDC-treated mice. (A-F) Values for DDC-treated mice only are shown here. Statistical significance of deviation from untreated control mice for each mouse strain is shown as green (upregulation compared to control) and orange (downregulation compared to control) asterisks above each group. Flow cytometric quantification of (A) Mo-MF (monocytes-macrophages, CD45^+^, CD11c^-^, CD11b^+^, Ly6C_6G^-^,F4/80^+^), (B) Gr-1^+^ cells (CD45^+^, CD11c^-^, CD11b^+^, Ly6c_6G^+^), (C) CD4^+^ T cells (CD45^+^, CD4^+^) and (D) CD8^+^ T cells (CD45^+^, CD8^+^) in liver tissue of untreated and 4 week DDC-treated mice of the indicated genotypes. Data from 4 week DDC-treated mice are graphically depicted. Wild type (WT) mice (untreated: n = 9, DDC-treated: n = 10), β7 integrin-deficient mice (β7^-/-^) (untreated: n = 9, DDC-treated: n = 10), L-selectin-deficient (L-sel^-/-^) (untreated: n = 10, DDC-treated: n = 8) and L-selectin/β7 integrin double-deficient (L-sel^-/-^/β7^-/-^) (untreated: n = 8, DDC-treated: n =8). Cells are shown as absolute numbers per gram liver. (E) Representative microscope images of multiplex immunofluorescence staining on paraffin sections of the liver. (F) Quantification of CD8^+^ T cells per mm^2^ of liver tissue. WT (untreated: n = 5, DDC-treated: n = 5), β7^-/-^ (untreated: n = 5, DDC-treated: n = 5) and L-Sel^-/-^ / β7^-/-^ (untreated: n = 5, DDC-treated: n = 5). Statistical significance was calculated by the unpaired t-test. Values are represented as mean ±SD and each dot (in a, b, c, d, and f) represents data from one mouse (* p < 0.05, ** p < 0.01, *** p < 0.001, ***p < 0.0001). (G) Ratio (Ro/Ri) of β7^-/-^ and WT CD8^+^ T cells isolated from the indicated organs 16 hours after adoptive transfer of an equal number of differently stained leukocytes of the respective genotypes into the lateral tail vein of WT recipients (n = 28-29). Statistical significance was calculated by one-sample t-test vs. 1 (* p < 0.05, *** p < 0.001).

### β7 integrin-expressing CD8^+^ T cells promote DDC-induced pathogenesis

To address the role of β7 integrin-expressing CD8⁺ T cells in DDC-mediated cholangiopathy cell transfer experiments were performed. To this end, we administered CD8^+^ T cells from WT mice intravenously (i.v.) into β7 integrin-deficient mice on the first day of a 4-week DDC treatment (Figure 4A). The subsequent development of cholangitis was assessed in comparison with that found in DDC-treated β7 integrin-deficient mice that had received an equal number of CD8^+^ T cells from β7 integrin-deficient mice. Transfer of CD8^+^ WT T cells significantly increased DDC-induced cholestasis in β7 integrin-deficient mice, as evidenced by decreased weight gain and increased serum levels of direct bilirubin and alkaline phosphatase (Figure 4B-D). In addition, it caused more severe cholangitis, which was reflected in a significantly increased porphyrin deposition and Sirius red stained fibrotic areas (Figure 4E and F), thereby restoring at least part of the susceptibility seen in WT mice. β7 integrin-deficient mice that had received an equal number of CD8^+^ T cells from β7 integrin-deficient mice were significantly less affected in all of the aforementioned parameters by the DDC-induced cholangiopathy.

**Figure 4:**
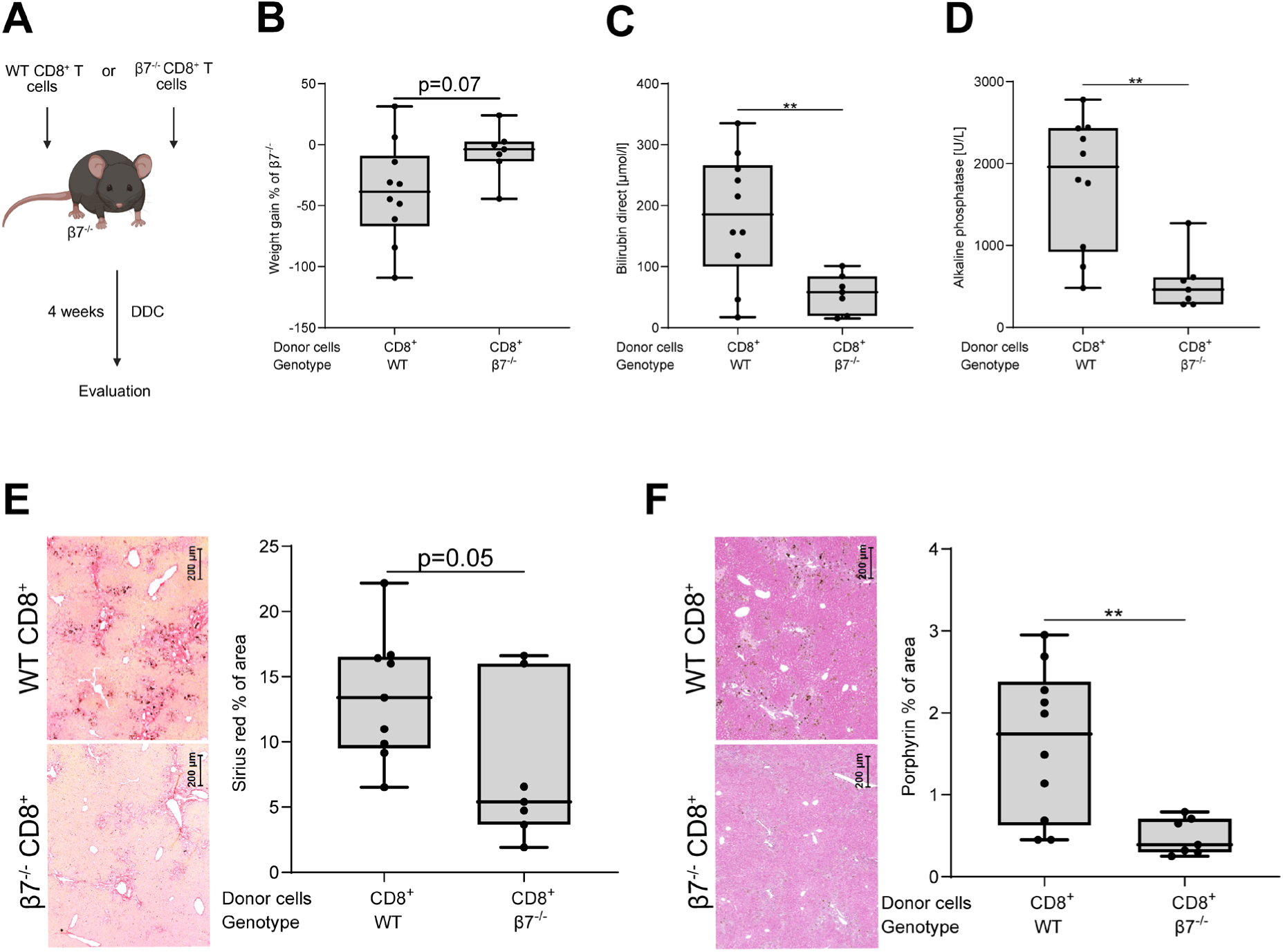
β7 integrin-expressing CD8^+^ T cells contribute to DDC-induced pathogenesis. (A) Experimental outline: β7 integrin-deficient (β7^-/-^) mice were adoptively transferred with CD8^+^ T cells from WT (n = 10) or β7^-/-^ (n = 7) mice and immediately subjected to DDC feeding. 4 weeks later serum and liver specimens were sampled and analyzed. (B) Percentage (%) of body weight gain. (C) Quantification of direct bilirubin and (D) quantification of Alkaline phosphatase in serum. (E) Representative photomicrographs of Sirius red stained liver sections from the indicated recipients at original magnification x10 and quantification of Sirius red stained collagen shown as percentage of the section area. (F) Representative photomicrographs of H&E stained liver sections from the indicated recipients at original magnification x10 and quantification of porphyrin deposition shown as percentage of the section area. Statistical significance was calculated by the unpaired t-test. Values are represented as mean ±SD and each dot represents data from one mouse (** p < 0.01).

### Amelioration of DDC-induced liver injury in adhesion molecule-deficient mice correlates with higher microbiota diversity and richness, and an improved intestinal barrier

Gut microbiota, intestinal barrier and bacterial-derived products seem to be involved in the pathogenesis of cholangiopathies and several studies have confirmed changes in the intestinal microbiota of PSC patients [25, 40, 41]. We used high-throughput 16S rRNA gene amplicon sequencing to investigate whether DDC altered the gut microbiota in WT mice and what influence adhesion molecule deficiency might have. Differences in fecal microbiota profiles between the genotypes before and after 4 weeks of DDC treatment were assessed by non-metric multidimensional scaling (NMDS) of generalized UniFrac distances and revealed drastic shifts due to DDC feeding with distinct profiles for each mouse strain (Figure 5A). Moreover, the species richness was significantly higher in feces of these mice strains after DDC treatment, but not when comparing the respective healthy mice strains (Figure 5B). In addition, fecal microbiota from DDC-treated β7 integrin-deficient and L-selectin/β7 integrin double-deficient mice showed increased Shannon effective counts compared to WT mice, a difference which was already seen in feces from untreated animals (Figure 5C). The relative abundance of *Bacteroidota* was substantially higher in feces of DDC-treated L-sel^-/-^ and L-sel^-/-^/β7^-/-^ mice in comparison to WT mice (Figure 5D), whereas the relative abundance of fecal *Bacillota* tended to be decreased (Figure 5E). The most pronounced DDC-induced difference at the family level across all genotypes was an increase in the relative abundance of *Prevotellaceae* (phylum *Bacteroidota*). Interestingly, the relative abundance of this family was significantly higher in feces of all adhesion molecule-deficient mice than in WT (Figure 5F). In addition, we observed a DDC-induced increase in the relative abundance of *Bacteroidaceae* (phylum *Bacteroidota*) in the feces of all mouse strains (Figure 5G) and a reduction in the relative abundance of *Muribaculaceae* (phylum *Bacteroidota*) (Figure 5H). Moreover, the relative abundance of *Lachnospiraceae* (phylum *Bacillota*) was significantly reduced in feces of all DDC-treated adhesion molecule-deficient mice in comparison to WT mice (Figure 5I).

**Figure 5:**
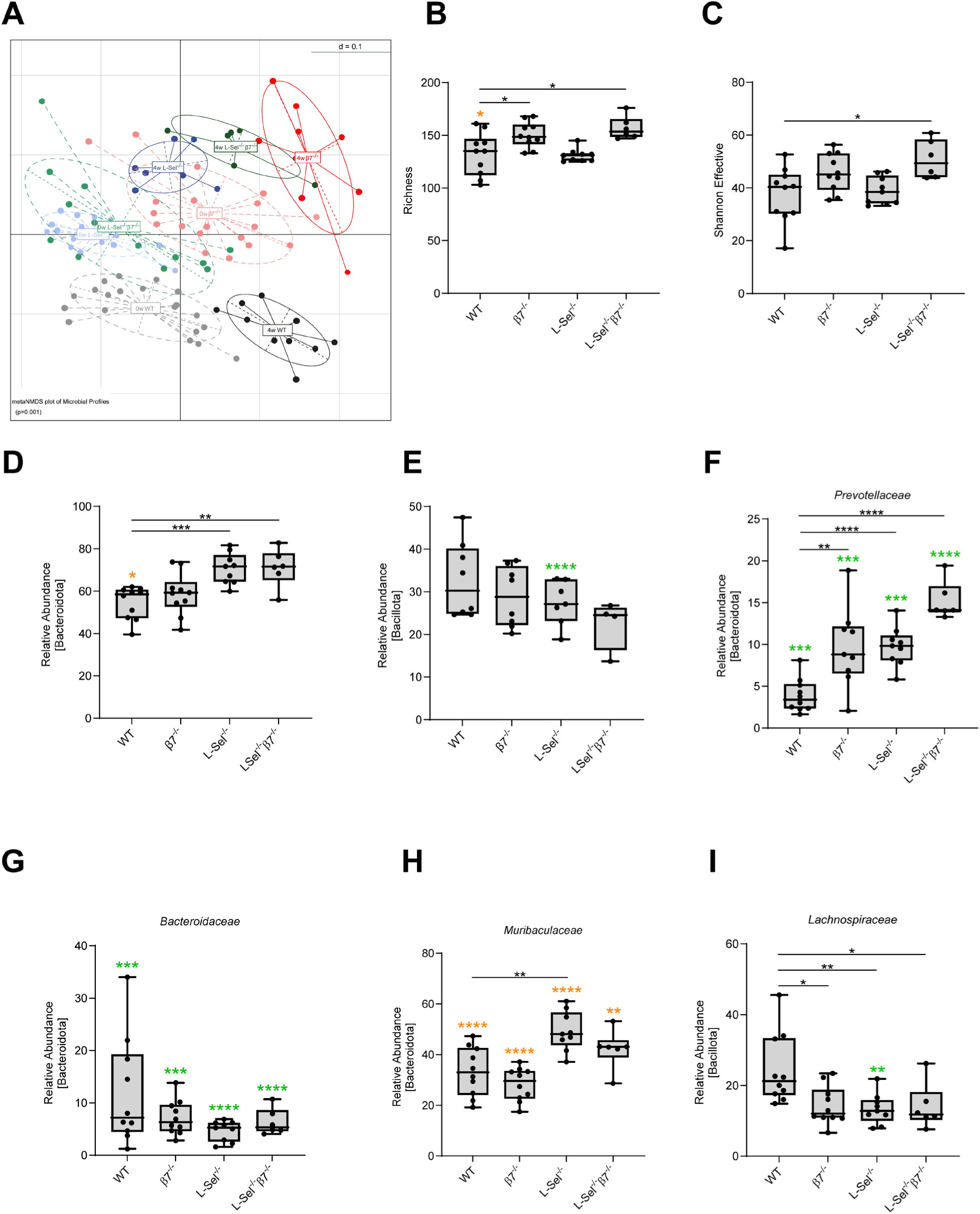
Changes in fecal microbiota in adhesion molecule-deficient mice. (A) metaNMDS plot of microbiota profiles based on generalized UniFrac distances from WT mice (untreated: n = 20, DDC-treated: n = 10), β7-integrin-deficient mice (β7^-/-^) (untreated: n = 19, DDC-treated: n = 10), L-selectin-deficient mice (L-sel^-/-^) (untreated: n = 17, DDC-treated: n = 9) and L-selectin/β7-integrin double-deficient mice (L-sel^-/-^/β7^-/-^) (untreated: n = 14, DDC-treated: n = 6). Each dot represents a mouse. (B-I) Values for DDC-treated mice only are shown here. Statistical significance of deviation from untreated control mice for each mouse strain is shown as green (upregulation compared to control) and orange (downregulation compared to control) asterisks above each group. Αlpha diversity as (B) richness and (C) Shannon effective counts. Relative abundance of (D) *Bacteroidata* and (E) *Bacillota* at the phylum level. Relative abundance of (F) *Prevotellaceae*, (G) *Bacteroidaceae*, (H) *Muribaculaceae*, and (I) *Lachnospiraceae*, at the family level. Statistical significance was calculated by the unpaired t-test. Values are represented as mean ±SD and each dot represents data from one mouse (*p < 0.05, ** p < 0.01, *** p < 0.001, ****p < 0.0001).

In addition to changes in the composition of intestinal microbiota, increased intestinal permeability has been described in cholangitis, supporting the hypothesis that translocation of bacterial products from the leaky gut into the liver contributes to liver inflammation [42]. We therefore evaluated mucosal permeability by measuring serum levels of FITC-dextran 4 hours post oral gavage. Our results demonstrate that DDC feeding is sufficient to increase intestinal permeability as DDC-fed WT and β7 integrin-deficient mice showed increased levels of FITC in serum. However, the concentration in serum of DDC-treated WT mice was significantly higher compared to β7 integrin-deficient mice, pointing to an improved gut barrier integrity in β7 integrin-deficient mice (Figure 6A). Although WT mice showed a slightly higher concentration of serum FITC-dextran even without a DDC diet compared to β7 integrin-deficient animals, this difference was not significant. In addition, we screened for signs of DDC-induced intestinal inflammation. None of the DDC-treated mouse strains showed apparent colitis. There was no difference in histology score for colitis, as evaluated blinded on H&E stained colon sections by a specialist histopathologist (Figure 6B and C), or for colon length (Figure 6D). Expression of several adhesion molecules correlates with disease activity of IBD [43]. Concerning the expression of adhesion molecules in the small intestine of DDC-treated mice, we observed a decrease in the mRNA level of *Icam-1*, irrespective of the genotype, which was less pronounced in L-selectin-deficient mice. The expression of *Madcam-1* and *Vcam-1* was significantly downregulated in the small intestine of DDC-treated β7 integrin-deficient mice when compared to respectively treated WT mice but was otherwise unchanged (Figure 6E). This finding correlates with the slightly lower inflammatory histology score of small intestines from β7 integrin-deficient mice (Figure 6F and G).

**Figure 6:**
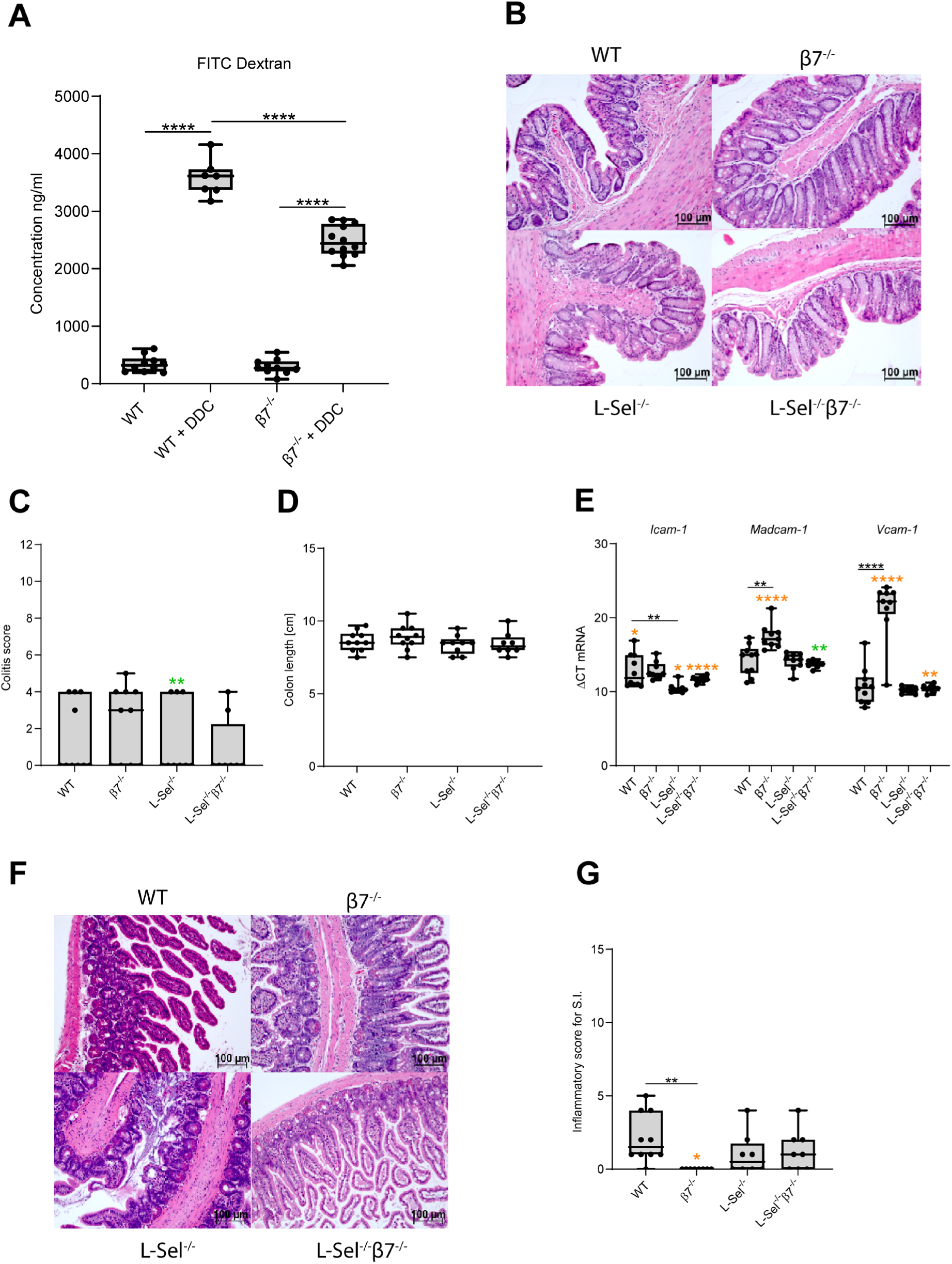
β7 integrin deficiency ameliorates DDC-induced gut barrier damage. (A) Analysis of the gut barrier from untreated and 4-week DDC-treated WT mice (untreated: n = 10, DDC-treated: n = 7) and β7-integrin-deficient mice (β7^-/-^) (untreated: n = 10, DDC-treated: n = 12) was measured by the FITC-dextran permeability assay. Serum concentrations of FITC-dextran were measured 4 hours after oral gavage. (B-G) Images and values for DDC-treated mice only are shown here. Statistical significance of deviation from untreated control mice for each mouse strain is shown as green (upregulation compared to control) and orange (downregulation compared to control) asterisks above each group. (B) Representative photomicrographs of H&E stained colon sections at original magnification x 20. (C) Evaluation of inflammatory score for Colitis (described in [44]). (D) Colon length. (E) Analysis of mRNA from specimens of the small intestine, that were sampled from the indicated untreated and 4-week DDC-treated mouse strains, by RT-PCR. Data from 4-week DDC-treated mice are graphically depicted. WT mice (untreated: n = 8, DDC-treated: n = 10), β7 integrin-deficient mice (β7^-/-^) (untreated: n = 9, DDC-treated: n = 9), L-selectin-deficient (L-sel^-/-^) (untreated: n = 10, DDC-treated: n = 9) and L-selectin/β7 integrin double-deficient (L-sel^-/-^/β7^-/-^) (untreated: n = 8, DDC-treated: n =8). For quantification, values are expressed as delta (Δ) ct values between the genes of interest and the housekeeping gene *β actin*. *Icam-1* (*intercellular adhesion molecule-1*), *Madcam-1* (*mucosal addressin cell-adhesion molecule-1*), *Vcam-1* (*vascular cell adhesion protein-1*). (F) Representative photomicrographs of H&E stained small intestine (SI) sections at original magnification x 20. (G) Evaluation of inflammatory score for S.I. (described in [45]). Statistical differences between treated groups were calculated by the unpaired t-test. Values are represented as mean ±SD and each dot (in a, c, d, and e) represents data from one mouse (*p < 0.05, ** p < 0.01, ****p < 0.0001).

## Discussion

A dysregulation of the complex interactions between the liver and the intestine and its enteric microbiota, leading to imbalanced inflammatory signals and increased immigration of inflammatory immune cells into the liver, seems to contribute to PSC development. The fact that IBD and PSC usually occur simultaneously led to the hypothesis that gut-homing markers expressing inflammatory lymphocytes are guided to the liver by the aberrant expression of adhesion molecules like MAdCAM-1, thereby contributing to inflammation in liver and gut [12–14]. To elucidate the role of the MAdCAM-1 binding partners α4β7 integrin and L-selectin in promoting PSC, we performed experiments in the well-established mouse model of DDC-induced cholangiopathy.

In line with earlier studies, DDC-feeding induced formation of protoporphyrin plaques resulting in pronounced biliary fibrosis affecting the small intrahepatic bile ducts [46]. In addition, we observed increased expression of MAdCAM-1, VCAM-1 and ICAM-1 in the liver, pointing to an inflammatory activation status of the liver endothelium in this model and the involvement of adhesion molecules in disease progression. Increased expression of gut-specific adhesion molecules including MAdCAM-1 has previously been reported in the livers of PSC patients and those with other chronic liver diseases [12, 13, 47, 48]. Most importantly, we show that β7 integrin-deficient, L-selectin-deficient, and L-selectin/β7 integrin double-deficient mice are markedly protected from DDC-induced liver injuries, reflected by less severe weight loss, reduced serum levels of liver injury markers, less liver fibrosis, and a reduced expression of bile metabolism-regulating proteins when compared to similarly treated WT mice. This finding clearly demonstrates the importance of these adhesion molecules in promoting PSC pathogenesis. A synergistic function of β7 integrin and L-selectin by differentially mediating rolling and firm adhesion had already been demonstrated in seeding mesenteric lymph nodes with lymphocytes [49].

Immune cell infiltration into the liver is a characteristic feature of PSC. DDC treatment caused a strong migration of CD8^+^ T cells into the liver and adoptive transfer of CD8^+^ WT T cells into DDC-treated β7 integrin-deficient mice was sufficient to increase disease susceptibility, whereas transfer of β7^-/-^ CD8⁺ T cells resulted in substantially less liver damage. This results confirms the fundamental role of β7 integrin-expressing CD8^+^ T cells in disease pathogenesis and is consistent with previous reports showing α4β7 expressing CD8^+^ T cells in the livers of PSC patients [13, 50] and that gut-primed CD8^+^ T cells were able to induce experimental immune-mediated cholangitis in an antigen-dependent manner [39]. Furthermore, the disease-promoting role of CD8^+^ T cells was recently described also in the PSC model of the mdr2 deficient mouse, where depletion of CD8^+^ T cells led to a significant attenuation of disease progression [38].

Unexpectedly, the increase of hepatic CD8⁺ T cells was even more pronounced in β7-integrin- and L-selectin/β7-integrin double-deficient mice compared to wild type controls and β7 integrin-deficient CD8^+^ T cells migrated significantly better into the liver than WT cells in short-term adoptive transfer experiments. This suggests that further adhesion molecules such as VCAM-1 or ICAM-1, which are also known to be elevated in PSC, contribute to hepatic recruitment. Moreover, increased numbers of hepatic CD8⁺ T cells in β7-deficient mice might result from the fact that in the absence of β7 integrin these cells are no longer retained in the gut. A comparable redistribution of lymphocytes has also been described in tumor models following downregulation of MAdCAM-1 [51, 52]

Increased migration of β7 integrin-deficient CD8^+^ into the liver, surprisingly, did not translate into increased disease activity suggesting that CD8^+^ T cells need β7 integrin to fulfill their pathogenic function. This hypothesis should be further investigated. It is supported by reports showing that E-cadherin binding is required to induce the cytolytic activity of CD8⁺ T cells [53] and that in PBC patients, a CD103⁺CD69⁺ CD8⁺ T-cell subset was found to infiltrate biliary epithelial cells via E-cadherin-mediated interactions [54].

Since the sole transfer of CD8^+^ T cells into DDC-treated β7 integrin-deficient mice did not reconstitute liver damage to the same degree as in correspondingly treated WT mice, it can be assumed that in addition to these, other immune cell types contribute to the pro-inflammatory action of β7 integrin in DDC-induced cholangitis. In this regard, it has recently been shown that inhibition of α4β7 integrin-mediated recruitment of CD4^+^ T cells by a blocking antibody reduces liver inflammation and fibrosis in a model of western-style induced MASLD and that β7 integrin-expressing CD4^+^ T cells contribute to Concanavalin A-induced acute immune-mediated hepatitis in mice [55, 56]. Moreover, recruitment of macrophages via the CCR2/MCP-1 axis has been found to promote pathogenesis in acute and chronic mouse models of sclerosing cholangitis [57] and Clodronate-mediated macrophage depletion to ameliorate DDC-induced liver damage [33]. Since DDC-treated β7 integrin-deficient mice tend to have reduced numbers of hepatic macrophages and significantly fewer CD4^+^ T cells compared to WT mice, it is quite reasonable to assume that a lack of these cells also contributed to the reduced disease activity.

Gut microbiota changes can affect inflammatory processes in the liver through activation of innate immunity, gut hormone production, and bile acid metabolism. Several studies have demonstrated such changes in PSC patients, pointing to a pathogenic role of gut microbiota in disease development [40, 41, 58–60]. Strategies targeting microbiota could therefore become useful therapeutic options. The potential influence of the gut microbiota on the expression of adhesion molecules, for instance through modulation of the availability of retinoic acid and vice versa, might be of special interest [27–30]. Independent of the genetic background of the mice investigated, DDC-feeding caused drastic shifts in the fecal microbiota profiles demonstrating that the microbiota in mice with DDC-induced cholangiopathy differed significantly from those of healthy control mice and might in part have caused the disease and contributed to its severity.

DDC treatment was accompanied by a decrease in the relative abundance of fecal *Bacteroidota*, which was less pronounced in adhesion molecule-deficient mice. This finding fits with reports of reduced gut *Bacteroidota* in patients with MASLD, metabolic dysfunction-associated steatohepatitis, liver cirrhosis, alcohol-associated liver disease, primary biliary cholangitis, and PSC [61, 62]. Within the *Bacteroidota*, we observed a DDC-induced increase in *Bacteroidaceae* and *Prevotellaceae*, while the relative abundance of *Muribaculaceae* was reduced. Several studies have reported decreased fecal abundance of the *Prevotella* in PSC patients, and *Segatella copri* (previously *Prevotella copri*) supplementation has been shown to ameliorate liver fibrosis in the DDC-induced PSC model [24, 40, 63, 64], suggesting a beneficial role of these bacteria for PSC pathogenesis. Although these findings are not consistent with the general DDC-induced increase in the relative abundance of fecal *Prevotellaceae* in our model, the strongly increased relative abundance observed in the adhesion molecule-deficient mice when compared to WT may have contributed to the improved disease outcome. Interestingly, the relative abundance of *Muribaculaceae* was strongly reduced in the DDC-induced cholangitis model. This finding is consistent with results from microbiota analysis in IBD, Type 2 diabetes, obesity, and patients with PBC and PSC, also showing a reduction in the occurrence of this family [65–67]. Overall, *Muribaculaceae* seem to play a beneficial role in maintaining host health and regulating the gut barrier [67], thus their absence might have had a negative impact on PSC pathogenesis.

The DDC–induced increase in the relative abundance of fecal *Bacillota*, which tended to be less pronounced in feces from adhesion molecule-deficient mice, is in line with reports showing increases in members of the phylum *Bacillota*, namely *Enterococcus*, *Lactobacillus*, and *Veillonella*, in the gut microbiota of PSC patients [23, 68]. The observed increase and higher relative abundance of *Lachnospiraceae* in the fecal microbiota of DDC-treated WT mice was not expected, as the SCFA producing *Lachnospiraceae* are thought to play a beneficial role, e.g. by driving Treg accumulation and differentiation, and negatively correlate with Mayo risk score [69, 70]. However, increased *Lachnospiraceae* have also been found in PSC-IBD patients and in feces of Mdr2-deficient mice, which develop cholangiopathy due to the absence of phospholipid from bile, while fecal microbiota transfer from these mice promoted liver disease in WT [63, 71]. Moreover, other species within the *Lachnospiraceae*, like *Ruminococcus gnavus*, are thought to promote mucin degradation and gut permeability and are positively correlated to Crohn’s disease [72, 73]. In conclusion, the observed DDC-induced changes in the overall gut microbiota, which differed significantly between the genotypes, might in part have been a cause of disease and also affected its severity.

As previously stated, enhanced gut permeability allowing unwanted translocation of products can trigger inflammatory responses in the liver and has been reported in PSC patients [22]. DDC-induced PSC was accompanied by a pronounced increase in intestinal permeability, which was significantly higher in WT than in β7 integrin-deficient mice. Unexpectedly, DDC treatment did not cause obvious histological intestinal inflammation, but actually led to downregulation of adhesion molecule expression in the small intestine, which was lowest in β7 integrin-deficient mice, pointing to the known protective effect of a β7 integrin-deficiency in chronic inflammatory bowel disease in mice and humans [74, 75]. The reduced basal inflammatory level and improved intestinal barrier integrity in β7 integrin-deficient mice might therefore have contributed to the observed protection from DDC-induced cholangitis.

In summary, we have shown that the MAdCAM-1 binding partners β7 integrin and L-selectin promote DDC-induced cholangitis, and that β7-integrin expressing CD8^+^ T cells play a decisive role in its pathogenesis. Moreover, by causing unfavorable microbiota shifts and destabilizing the gut barrier, adhesion molecule-dependent interactions of immune cells might contribute to the inflammatory process. These findings further our understanding of the processes underlying the development of cholangiopathies, which will be important for the development of new therapies.

## Materials and Methods

### Ethical Statement

The study was carried out in accordance with the regulations laid down by the regional authorities for nature, environmental, and consumer protection of North Rhine-Westphalia (LANUV, Recklinghausen, Germany) and approved by the respective Committee (Permit Number: 81-02.04.2017.A429 and 81-02.04-2021.A402). All experiments were performed in accordance with the German guidelines for animal housing and husbandry. Treatments fulfilled the criteria of the German administrative panel on laboratory animal care.

### Housing, mice, and dietary treatments

Animals were housed in specific pathogen-free conditions with 12 h light/dark cycles and water and food available ad libitum. As PSC affects men more commonly than women [76] and to reduce variability due to the menstrual cycle and thereby limit mouse numbers, all experiments were performed with 10–14-week-old male β7-integrin-deficient (β7^-/-^) (C57BL/6-Itgb7tm1Cgn/J)[16], L-selectin-deficient (L-sel^-/-^) (B6;129S2-Selltm1Hyn/J)[77], L-selectin/β7-integrin double-deficient (L-sel^-/-^/β7^-/-^) (B6;129S2-Selltm1Hyn C57BL/6-Itgb7tm1Cgn/J)[49], and wild type (WT) mice on C57BL/6J background from the same barrier and room. Mice were fed chow (9 kcal % fat, 24 kcal % protein, 67 kcal % carbohydrates) (ssniff, Soest, Germany, rat/mouse–maintenance) or a 3,5-diethoxycarbonyl-1.4-dihydrocollidine (1g/kg) containing diet (Altromin International, Lage, Germany). All experiments were repeated in at least two independent experimental setups.

### Determination of serum parameters

Bilirubin, serum aspartate aminotransferase (AST), serum alanine aminotransferase (ALT) and alkaline phosphatase (ALP) in serum were measured by the Central Laboratory Facility of the University Hospital, RWTH Aachen.

### Histological stainings and evaluation

4 μm formalin-fixed paraffin sections of liver and gut were stained with the H&E staining kit or Picro Sirius red Solution (Abcam, Cambridge, England) in accordance with the manufacturer’s instructions. Histopathological scoring of sections from colon (maximum achievable score 12) and small intestine (maximum achievable score 15) and their validation was performed blinded via a colon or small intestine score, as described previously [44, 45].

### Multiplex immunfluorescence staining

Multiplex immunofluorescence staining on 2 μm thick, formalin-fixed paraffin sections of the liver and analyses were performed as previously described [78, 79]. To restore the immunoreactivity of the paraffin-embedded tissue, antigen retrieval was performed using a universal HIER antigen retrieval reagent. This was followed by staining with the antibody and counterstaining with 4′,6-diamidino-2-phenylindole (DAPI). The antibodies used in this study are listed in Table 1.

**Table 1:** Antibodies for multiplex immunofluorescence staining.

| 1. Antibody | Host | Dilution | Manufacturer | Order number |
| --- | --- | --- | --- | --- |
| CD8a | Rat | 1:200 | Thermo Fisher Scientific GmbH | 14-0808-82 |
| Cytokeratin 7 | Rabbit | 1:500 | Abcam | ab181598 |
| 2. Antibody | Host | Dilution | Manufacturer | Order number |
| Anti-rat IgG (H+L), (Alexa Fluor® 647 Conjugate) | Goat | 1:500 | Cell Signaling Technology | 4418S |
| Anti-Rabbit IgG (H+L) Cross-Adsorbed Secondary Antibody, Alexa Fluor™ 750 | Goat | 1:500 | Thermo Fisher Scientific GmbH | # A-21039 |

The multiplex immunofluorescence staining images were taken using a Zeiss AxioObserver 7 (Carl Zeiss Microscopy, Oberkochen, Germany). All other images were acquired using an Axioplan2 microscope (Carl Zeiss Microscopy, Oberkochen, Germany) and Zen lite 3.2 software (Carl Zeiss Microscopy, Oberkochen, Germany).

### Flow cytometry

Liver cell isolation and surface stainings were performed as described previously using combinations of the monoclonal antibodies listed in Table 2 [80] and quantified using the gating strategy shown below (Figure 7). All flow cytometric measurements were performed on a Canto-II cytometer (BD Biosciences). Data were analyzed by FlowJo 8.7.2 and 10.2 software (Tree Star, Ashland, OR, USA).

**Figure 7:**
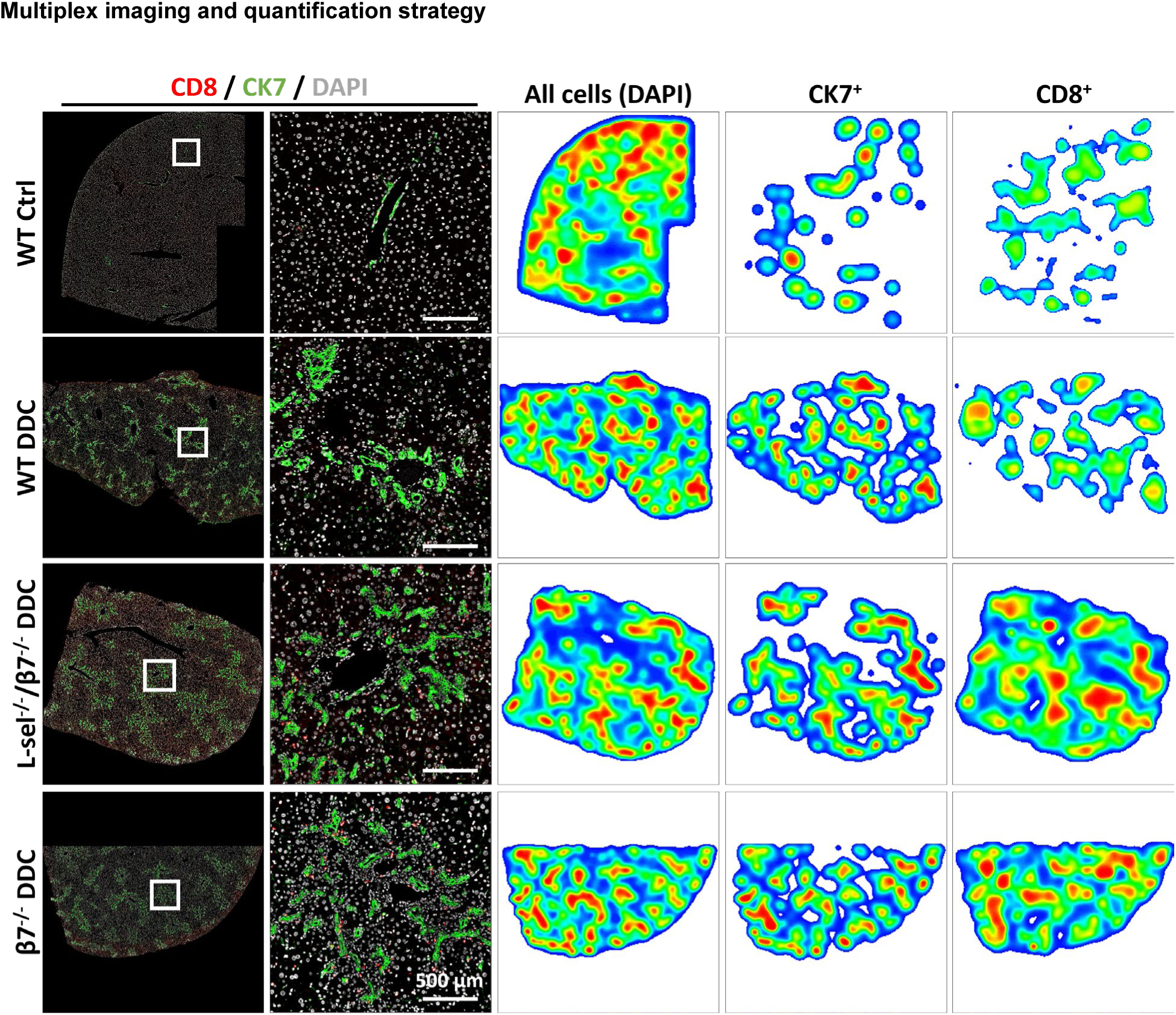
Distribution of CD8⁺ T cells in liver tissue. Representative immunofluorescence images of liver sections stained for CD8 (red), CK7 (bile ducts, green), and DAPI (nuclei, white). Enlarged views of the indicated regions (white boxes) are shown in the second column. Quantitative heatmaps of CD8⁺ T cell density are displayed in the right panels, illustrating regional clustering around CK7⁺ bile ducts. Scale bar: 500 µm.

**Table 2:** Antibodies used in this study.

| Antibody | Channel | Manufacturer | Clone | Dilution |
| --- | --- | --- | --- | --- |
| CD4 | APC | BD Biosciences | RM4-5 | 1:200 |
| CD45 | APC-Cy7 | BD Biosciences | 30-F11 | 1:400 |
| CD8a | AmCyan | BioLegend | 53-6-7 | 1:400 |
| CD11b | Amcyan | BD Biosciences | M1/70 | 1:400 |
| CD11c | APC | eBioscience | N418 | 1:200 |
| F4/80 | eFluor 450 | Serotec | Cl:A3-1 | 1:200 |
| Ly6G/6C | FITC | eBioscience | R86-8C5 | 1:200 |
| CD4 | PE | BD Biosciences | GK1.5 | 1:400 |
| CD3 | PerCP-Cyanine5.5 | eBioscience | 145-2C11 | 1:100 |
| CD19 | PE-Cy7 | BioLegend | 1D3 | 1:200 |
| NK 1.1 | FITC | BioLegend | PK136 | 1:600 |
| Calibrite counting beads | APC | BD Biosciences |  | 1 million beads/ml PBS |

### Measurement of intestinal permeability

Mice were fasted for 4 hours while maintaining ad libitum access to water. Then fluorescein isothiocyanate (FITC)-dextran 4 (FD4) (TdB Labs AB) was administered by oral gavage at a dose of 0.6 mg/g body weight and 4 hours later serum was collected. The fluorescence intensities of duplicates from 1:1 diluted serum samples and a standard curve were measured fluorometrically (excitation 485 nm, emission 528 nm; SpectraMax iD3 from Molecular Devices) and the concentration of FITC dextran in mouse serum was calculated.

### Short term migration assay

Pooled single-cell suspensions of mesenteric lymph nodes, peripheral lymph nodes, brachial lymph nodes and spleen from WT and β7^-/-^ mice were labeled at 1:4000 dilution with either cell proliferation dye eFluor 450 or e677 according to the manufacturers’ instructions (eBioscience, Frankfurt am Main, Germany). eFluor 450-labeled leukocytes were mixed with an equal number of e677-labeled leukocytes and injected (2-3 x 10^7^ total cells) in a volume of 100µl into the lateral tail vein of WT recipients. Sixteen hours later, ratio of eFluor 450-e677-labeled CD8^+^ T cells was determined by flow cytometry in single-cell suspensions isolated from liver, spleen and Peyer’s patches of recipient mice. To correct for the input ratio of injected cells, an aliquot of the injections mixtures was analyszed for the ratio of eFluor 450-e677-labeled cells (Ri).

### Lymphocyte isolation and adoptive cell transfer of CD8^+^ T cells

Single cell suspensions from spleen, MLN, and PLN were pooled after erythrocyte lysis with lysis buffer (BD Pharmingen, San Jose, CA). CD8^+^ T cells purification was performed by magnetic-activated cell sorting (MACS) using the CD8a T-cell isolation kit for negative selection from Miltenyi (Miltenyi Biotec, Bergisch Gladbach, Germany) according to the manufacturer’s instructions. Purity of the isolated cell fractions was controlled by flow cytometry. The cells were resuspended in an appropriate volume of sterile phosphate-buffered saline allowing for 2 x 10^6^ cells per tail vein injection and were transferred once on the first day of DDC treatment.

### 16S rRNA gene sequencing

Fecal samples were collected from co-housed DDC-fed WT, β7 integrin-deficient, L-selectin-deficient and L-selectin/β7 integrin double-deficient mice at the start (0 weeks) and end point of feeding (4 weeks). Metagenomic DNA was extracted from frozen fecal pellets by bacterial disruption using zirconia beads (Biospec) and a FastPrep 24 device (MP Biomedical, USA) at 6 m/s for 20 s. Homogenates were incubated at 80 °C for 10 min and centrifuged at 17000×g for 1 min. The supernatant was taken for DNA isolation using the QIAamp fast DNA stool kit (Qiagen, Cat. No. 51604) according to the supplier’s instruction. The hypervariable regions were used to create the Next Generation Sequencing [NGS] library V3/V4 of the 16S rRNA gene from bacteria in stool. 341F and 785R were used as universal primer binding sites. The purification took place through the AMPure XP system, followed by a subsequent quality control and a fluorometric DNA quantification. Sequencing was performed with the Illumina MiSeq DNA sequencer. The full protocol has been described by Lagkouvardos and colleagues [81].

As the amplicon of the V3/V4 region is approximately 464 base pairs, sequences shorter than 350 base pairs and longer than 500 base pairs were excluded from the analysis. The ends of both sequences were shortened by 15 base pairs each and the expected error was 2. Sequencing depth was evaluated by means of rarefaction curves and, as apparent from the observed plateau in these curves all samples passed the quality check and were analyzed further (Figure 8). The sequences obtained were transferred to the "integrated microbial NGS platform [imngs]" program. To classify groups of closely related bacteria, operational taxonomic units (OTUs) were generated with 97 % agreement by the USEARCH 11 program. Across the 105 samples studied, a total of 2,936,451 high-quality and chimera-checked sequences (16,779±15,976) were analyzed, representing 294 operational taxonomic units (OTUs) (141±18 OTUs per sample). There was a filter for the abundance of the microbes, which was >0.25 %. Sequence alignment and taxonomic classification were conducted with SINA 1.6.1, using the taxonomy of SILVA release 128 [82]. Downstream analysis was performed in the R programming environment using Rhea (https://lagkouvardos.github.io/Rhea/) [83]. OTU tables were normalized to account for differences in sequence depth. Βeta-diversity was computed from generalized UniFrac distances [84]. Αlpha-diversity was assessed based on species richness and Shannon effective diversity [85].

**Figure 8:**
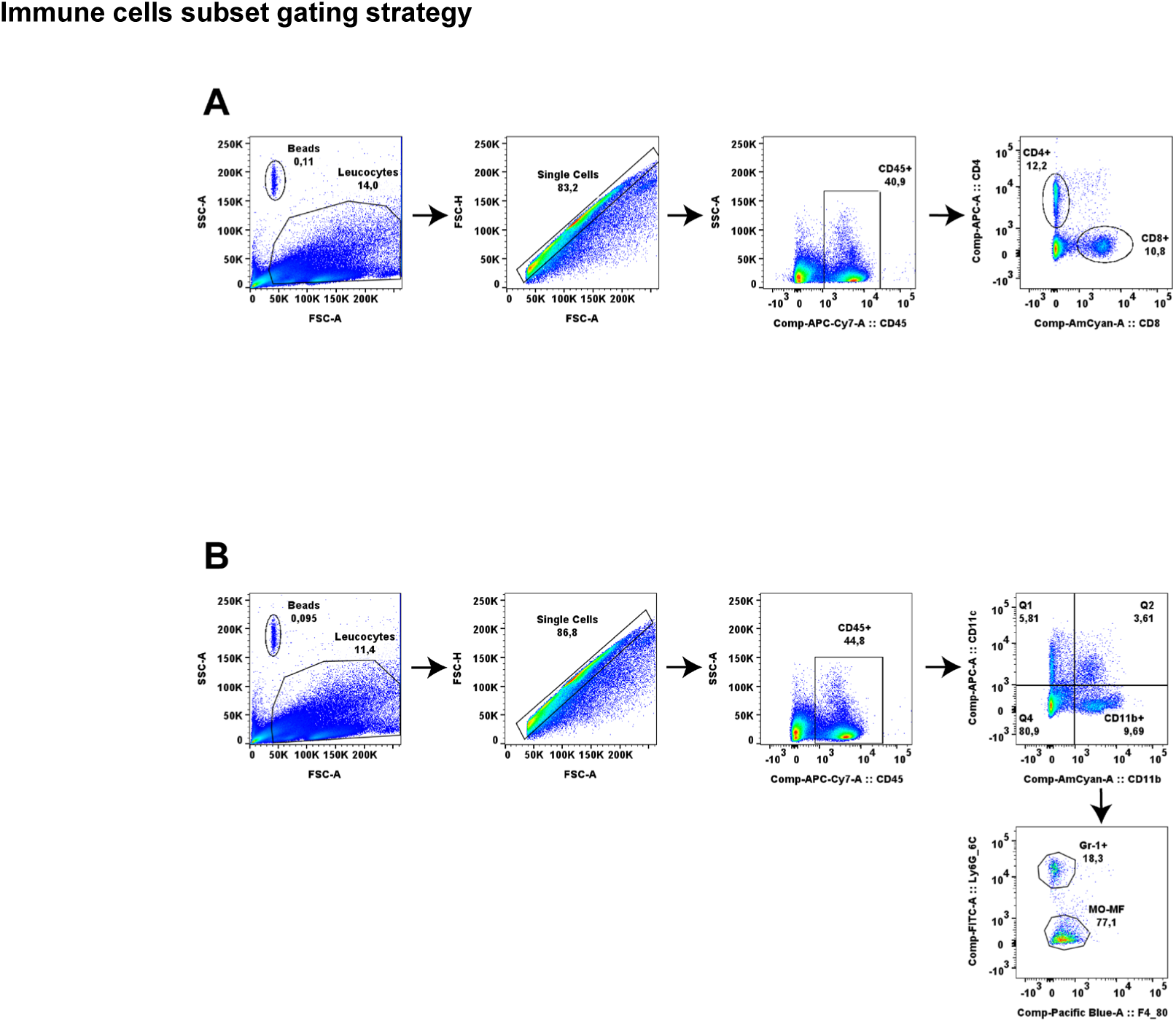
Immune cell subset gating strategies for liver cell profiling by multiparameter flow cytometry. (A) Gating for lymphoid cells including CD4^+^ T cells (CD45^+^ CD4^+^), CD8^+^ T cells (CD45^+^ CD8^+^) and (B) Gating for myeloid cells including monocytes-macrophages (Mo-MF) (CD45^+^CD11c^-^, CD11b^+^, F4/80^+^), Gr-1+ cells (CD45^+^CD11c^-^CD11b^+^Ly6C_6G**^+^**).

**Figure 9:**
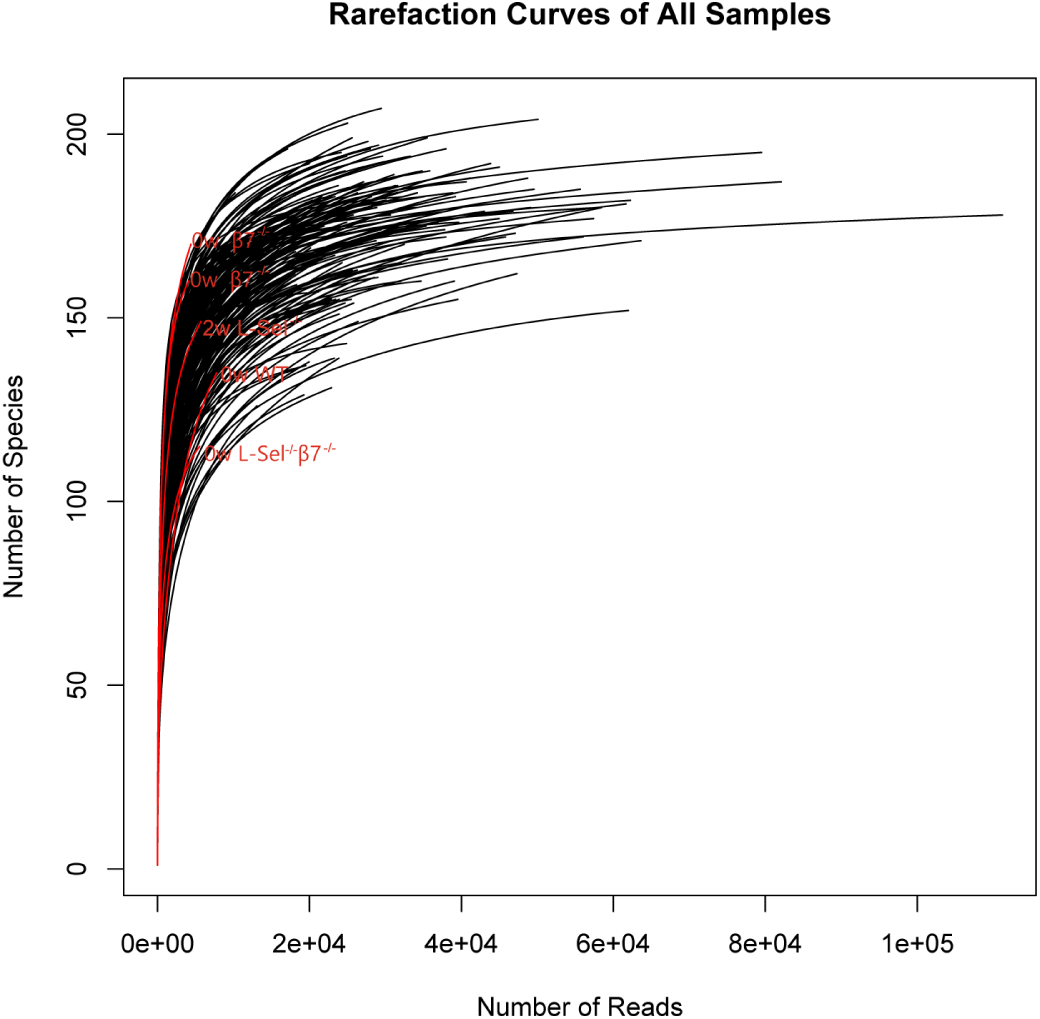
Rarefaction curves were constructed to illustrate sequencing depth. The number of molecular species observed in each sample was plotted against the number of reads acquired to ensure sufficient sequencing depth. The curve was generated in R using Rhea. The five samples with the least sequencing reads are color-coded in red.

### Real-time PCR

Total RNA isolations from the liver and complementary DNA (cDNA) synthesis were performed with the RNeasy Protect Mini Kit (Qiagen GmbH, Hilden, Germany) and Transcriptor First Strand cDNA Synthesis Kit (Roche Diagnostic GmbH, Mannheim, Germany) according to the manufacturer’s instructions. Real-time polymerase chain reactions (RT-PCR) were performed in duplicate using the quantitative (q)PCR Master Mix for SYBR Green I (Eurogentec, Cologne, Germany), in a total volume of 20 µL, on a 7300 RT-PCR system with 7000 System SDS Software Version 1.2.3 (Applied Bioscience, Darmstadt, Germany). Primer sequences are listed in Table 3 β *Actin* was used as endogenous control for normalization.

**Table 3:**
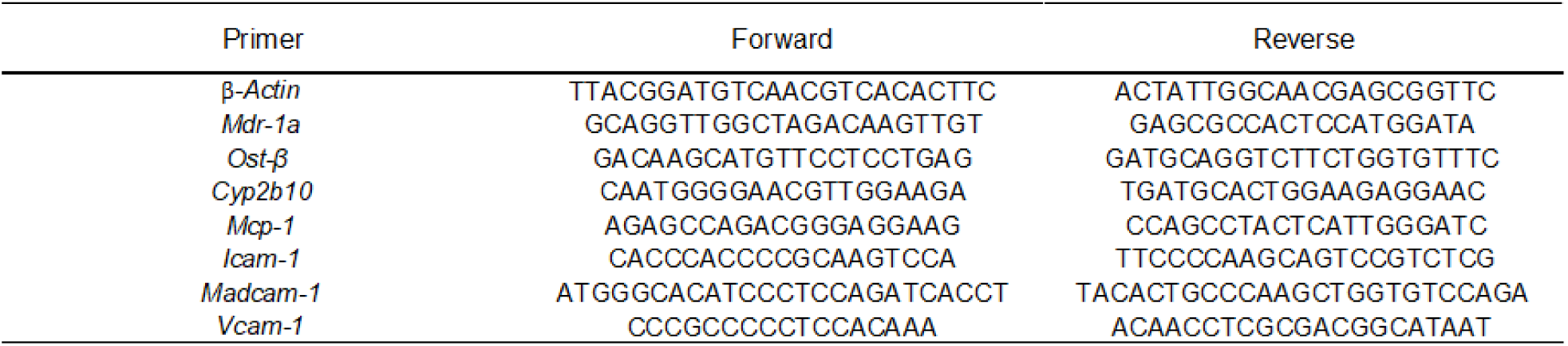
Primers used in this study.

### Statistical Analysis

Unless otherwise indicated, statistical analysis was performed with GraphPad Prism software (version 10 GraphPad, La Jolla, CA, USA). Data are presented as mean ± standard deviation (SD). Significance values were calculated using the Student’s t-test when comparing two groups or two-way analysis of variance (ANOVA) and Tukey post-test. Values of p < 0.05 were considered significant (* p < 0.05, ** p < 0.01, *** p < 0.001, and **** p ≤ 0.0001).

Statistical analysis of data from microbiota sequencing was performed in the R programming environment using Rhea (https://lagkouvardos.github.io/Rhea/)[83].

## Acknowledgments

This work was supported by the "Immunohistochemistry facility", a core facility of the Interdisciplinary Center for Clinical Research (IZKF) Aachen within the Faculty of Medicine at RWTH Aachen University

## Abbreviations

ALT: alanine aminotransferase;
ALP: alkaline phosphatase;
AST: aspartate aminotransferase;
CAR: constitutive androstane receptor;
DDC: 3,5-diethoxycarbonyl-1,4-dihydrocollidine;
FITC: Fluorescein isothiocyanate;
IBD: inflammatory bowel disease;
ICAM-1: intercellular adhesion molecule-1;
MACS: magnetic-activated cell sorting;
MAdCAM-1: mucosal addressin cell-adhesion molecule-1;
MASLD: metabolic dysfunction-associated steatotic liver disease;
MCP-1: monocyte chemoattractant protein-1;
MDR1: multidrug-resistance protein 1;
OST-β: organic solute transporter beta;
OTUs: operational taxonomic units;
PSC: primary sclerosing cholangitis;
RT-PCR: real-time polymerase chain reaction;
VCAM-1: vascular cell adhesion molecule-1;
WT: wild-type

## Data availability statement

The microbiota sequencing data presented in this study were submitted to the Sequence Read Archive and are openly available under the accession number PRJEB82426.

